# Supervised machine learning versus expert assessment of ultrastructural changes in wild-type and OGT knockout macrophages

**DOI:** 10.64898/2026.01.21.700795

**Authors:** Thijs van der Meer, Graham A. Heieis, Bart Everts, Frank G. A. Faas, Roman I. Koning

## Abstract

Automated transmission electron microscopy (TEM) generates large datasets that challenge traditional qualitative analysis of cellular ultrastructure. Quantitative assessment of structural differences between different samples remains difficult due to structural variability in thin sections of organelles. Here we applied supervised machine learning (sML) to segment, quantify and compare cellular structures -including nuclei, chromatin, mitochondria, rough endoplasmic reticulum, and endocytic vesicles- in large TEM images of wild-type macrophages versus those with altered cellular physiology due to deficiency in O-GlcNAc Transferase (OGT). sML revealed that OGT knockout macrophages are larger and more oval, with increased euchromatin, nucleoli size, and relative mitochondrial and rER surface areas. Comparison with six TEM experts showed sML provides more objective and sensitive quantification of subtle differences, while expert consensus is only achieved for larger structural variations. These findings demonstrate that sML enhances quantitative TEM analysis and complements human expertise in ultrastructural studies.

## Introduction

Over the last decade the pace at which microscopy images of biological structures are acquired has increased dramatically. As a consequence of the increase in number and size of the datasets, image processing of electron microscopy data is becoming increasingly important. In cryo electron microscopy (cryoEM) thousands of movies are automatically acquired from multiple samples for several days. Image processing is focused on image corrections, increasing signal-to-noise ratio of the noisy images, and generating three-dimensional structures from projected images. This results in significant data reduction, using single particle analysis (SPA) resulting in protein structures, and using cryo electron tomography (cryoET) resulting in a cellular volume reconstructions (Baldwin et al., 2018). In scanning electron microscopy (SEM) huge datasets are recorded using serial block face imaging or multibeam imaging. Image analysis In SEM is primarily focused on 3D segmentation of biologically significant structures, such as cell outlines and organelles, in the large image datasets, subsequent shape analysis and quantification (Titze & Genoud, 2016). Furthermore, in light microscopy (LM) automated whole slides imaging is routine and automation is 2D image analysis is well developed (Li et al., 2022).

While image processing for cryoEM, 3D SEM and 2D LM slides is becoming more routine, 2D image analysis of large TEM datasets remains less well advanced. Since the development of virtual nanoscopy transmission electron microscopy (TEM) imaging (Faas et al., 2012), we have routinely acquired large-scale nanometer-resolution virtual slides of plastic embedded thin sections from cells and tissues (Cabukusta et al., 2024; de Boeck et al., 2016; Giacomelli et al., 2020; Koning et al., 2022; Melia et al., 2017; Ravelli et al., 2013). Recently, we developed a software interface and workflow for supervised machine learning (sML) annotation, modelling, prediction, and visualization in composite image TEM maps. This approach enables segmentation of histologically relevant subcellular structures, such as nuclei, mitochondria, rough endoplasmic reticulum, lipid droplets, etc., and use these for quantification (manuscript in preparation).

TEM virtual slides are routinely used to analyze and compare histological differences between different samples. However, examination of large TEM slides is time consuming, human expert analysis is subjective, and it is difficult and compare subtle structural differences in large images. Even more, quantifying differences of the various biological structures between differences samples, e.g. knock out and wild type cells, is difficult, given the natural biological variation within one sample. In order to get a reproducible analysis of cellular structural, we investigated whether we could quantify variations of sML segmented structures between different samples.

For this case study, we compared murine macrophages that were deficient for O-linked β-N-acetylglucosamine (O-GlcNAc) transferase (OGT) with wild-type mouse macrophages. O-GlcNAc is an enzyme that post-translationally adds N-acetylglucosamine (N-GlcNAc) to threonine and serine residues in proteins. O-GlcNAcylation couples nutrient sensing, metabolism, signal transduction and transcription and is involved in many diseases, including neurodegenerative diseases, diabetes, cancers and immunology (Chang, Weng, & Lin, 2020). Initial light microscopy and biochemical experiments on macrophages from Lysm-cre* Ogt-flox mice (from hereon referred to as OGT KO macrophages) pointed towards profound changes in cellular physiology, including structural changes of the rER, mitochondria and Golgi, providing a highly suitable experimental setup to train and validate our sML model on(Heieis et al., 2026; Yang et al., 2020). We started out by manually annotating and making CNN models of whole macrophages, nuclei, euchromatin, heterochromatin, nucleoli, mitochondria, rough ER, Golgi stacks and endocytic vesicles. These models were used to predict these structures in virtual TEM slides using sML and the segmentations were used for quantitatively analyze the differences in occurrence and size between WT and OGT KO macrophages. Moreover, to validate sML, we compared the quantitative results with the human analysis of six TEM experts from our lab.

Structures segmentation of cellular components using sML, followed by quantification revealed large differences between OGT KO and WT mouse macrophages of cell size and shape, nuclear size, the distribution of heterochromatin and euchromatin, and number and shape of mitochondria. The observed changes in occurrence and size provided valuable insight into the underlying biochemical differences between WT and OGT KO macrophages. Comparison of sML results with TEM expert analysis, showed that sML quantification is in accordance with human analysis for large size differences, but revealed superior capacity to detect more subtle size differences and quantification differences.

## Materials and Methods

### Animals

For this study, Lyz2-cre and Itgax-cre mice were crossed with Ogt-flox mice and bred in-house under specific pathogen free (SPF) conditions. All mice were housed in a 12h:12h light-dark cycle at 25°C. The mice had access to food and water ad libitum. All animal experiments were approved by the Leiden University Animal Ethics Committee, as well as approved by the Dutch Central Authority for Scientific Procedures on Animals (CCD). Animal license number AVD1160020198846

### Sample preparation

The mice were sacrificed using CO_2_ euthanasia. Macrophages were harvested from the peritoneal cavity of both WT mice and OGT KO mice respectively (Zhang, Goncalves, & Mosser, 2008). In short, PBS was injected into the peritoneal cavity and sucked up again with the macrophages. To preserve the structure of the macrophages, the cells were fixated at a final concentration of 1.5% glutaraldehyde in 0.1M cacodylate buffer by adding double-concentrated fixative to the collected cells. To create a pellet, the cells were centrifuged at 2500 rpm for 10 minutes. The cells were rinsed with 0.1M cacodylate buffer, after which post-fixation was performed using osmium tetroxide (1% OsO4 and 1.5% potassium hexacyanoferrate(III) in MilliQ) on ice for 1 hour. The fixed cells were centrifuged at 4000 rpm for 5 minutes and again rinsed using 0.1M cacodylate buffer. Then, the fixative was resuspended in agar (2% bacto-agar in MilliQ). Again, the samples were centrifuged at 5000 rpm for 5 minutes. When the agar had set, the pellet was cut off, put in 70% ethanol (EtOH), and left overnight at 4°C. The next day, the cells were dehydrated using a series of 80% EtOH and 90% EtOH for 10 minutes each. At last, the samples were put in 100% absolute EtOH twice for 30 minutes. For embedding, the samples were put through a series of Epon/Acetone mixtures. Using subsequent Epon/Acetone ratios of 1:2, 1:1, and 2:1 each for 30 minutes, followed by an hour of 100% Epon, the samples were fully embedded. They were transferred to BEEM® capsules (Leadd) and left for two days at 70°C. When the samples were hardened, 90 nm or 45 nm thick slices were cut using a diamond knife on an ultra-microtome (Leica EM UC6). To confirm the presence of cells in semithin slices, a drop of toluidine blue was put on the first couple of slices, and this was checked with a light microscope. After the confirmation of the presence of cells on a slice, the rest of the sample was cut, and the slices were captured on a copper grid (mesh 50). To stain the cells, the copper grids were exposed to a droplet of 7% uranyl acetate for 10 minutes. Then, the grids were washed with 5 droplets of MilliQ and put on a droplet of 0.01M NaOH. The staining was finished by putting the grids on a droplet of lead citrate for 5 minutes followed by a washing step using 0.01M NaOH six times. The grids were air-dried and ready for imaging.

### Electron microscopy

The TEM images were acquired using an Tecnai 12 Spirit Twin transmission electron microscope (FEI) equipped with an Eagle 4k x 4k CCD camera (FEI). Images were taken at 6500x nominal magnification at 120 kV. An emission current between 0.5 µA and 2.5 μA and a defocus of 2 μm was used for all images. MyTEM and MyStitch software (Faas et al., 2012) was used to record consecutive images with a 20% overlap and stitch these into a composite virtual TEM slide. These virtual TEM slides (which we call TEM stitches) are composed of hundred to thousands of connecting CCD images and saved as a Tiff pyramid with different resolution levels.

### Supervised Machine Learning

Supervised machine learning is a method where the user provides annotated data that it used to train a numerical model. This model is used to predict similar structures in unseen data using a convolutional neural network (CNN). For our machine learning workflow, we used TEM stitches that were processed in a custom written Composite image Annotation, Visualization and Image Analysis (CAVIA) software via a browser interface (manuscript in preparation). In short, boxes were placed in a TEM stitch to define the regions used for neural network training. In these boxes, all structures for which a model needs to be trained are manually annotated. Boxes and annotations can be done at any level in the Tiff pyramid, i.e., the magnification, and predictions are also done at this level. Annotation, modelling and predictions were done at higher resolution for smaller structures (e.g. rough endoplasmic reticulum) and while larger structures (e.g. the nucleus) were processed at lower resolution levels to optimize model calculation and prediction times. Models were created using TensorFlow (Abadi, 2015) and Keras (Chollet, 2015). Typically, three rounds of annotation, modelling, and predictions (initial annotation, removal of false positives and addition of false negatives) were used to optimize the model. Minor manual corrections on the final predictions of the cell and nucleus were done prior to data quantification.

### Annotations

Manual annotations of structures were performed by a single master student without prior EM expertise using the following guidelines. Macrophages were distinguished from lymphocytes by the presence of endocytic vesicles (lymphocytes have little to no endocytic vesicles), and their size (macrophages are larger). Nuclei were distinguishable by their central cellular location, size and morphology. Dark nuclear areas were assigned as heterochromatin, whereas lighter nuclear areas were marked as euchromatin. An nuclear area was assigned as nucleolus when it was approximately circular in shape and had a grey level in-between those of the heterochromatin and euchromatin. Mitochondria were recognised by their double membrane and cristae. Rough endoplasmic reticulum (rER) was only annotated when these resembled rod-like membrane structure with ribosomes (small black dots). Any differently structured thin section of the rER, even when recognized as (part of the) rER, was not annotated to prevent too much variation and ambiguity of the model. From the Golgi apparatus only stacked membrane compartments were annotated since it was difficult to discriminate Golgi-derived vesicles from other types of vesicular structures and smooth ER. Both the Golgi apparatus and nucleoli were completely manually annotated and not modelled for sML as there were too few instances to generate a reliable model. It was therefore both faster and more accurate to manually annotate them all.

### Image analysis

Segmentations of predicted and manually annotated structures were extracted and their occurrence, size, roundness, and surface area (in percentage) were calculated with a custom Python 3.8.10 script using Spyder 5.5.3. For calculation of the surface area typically a primary segmentation (whole macrophages or nuclei) was chosen in which the presence of a secondary segmentation (nucleus, mitochondria, endocytic vesicles, rER and Golgi for macrophages; heterochromatin, euchromatin, and nucleoli for nuclei) was calculated. The occurrence represented the average number of secondary structures within a single primary structure. The size depicted the area of the segmented objects. The roundness enumerated the circularity of an object, between 1 (perfectly round) to more elongated (< 1). The surface area represented the relative total area of a structure (e.g. all mitochondria in all macrophages, or all nucleoli in all nuclei).

Since macrophages and nuclei were used as primary structures for the calculations, all predictions of the macrophages and nuclei were manually corrected to optimise the correctness of the quantifications. With the current set-up of the program, it was only possible to obtain data on structures that were inside another structure. For example, heterochromatin, euchromatin, and nucleoli data were only counted if present within the predicted area of the nuclei, ignoring all (false positive) predictions present outside the nuclei. Also, (incorrectly) predicted pixels that were present as single pixels or patches, smaller that a certain minimum size were occasionally removed.

### Expert analysis

To validate the statistical results and quantifications from the machine learning predictions, with more traditional observations, TEM experts were asked to examine all WT (n=4) and OGT KO (n=4) stitches and fill out a survey. In total six TEM experts completed the survey, 3 male, 3 female, aged between 46 and 73 years of age, all with practical TEM experience ranging between 20 to 45 years,. Virtual slides were accessed, without any time constraints, via different tabs in a web-based image browser. The experts did not have access to the annotations, except for the macrophage segmentations, since also lymphocytes were present in the samples.

## Results

### Image acquisition and segmentation

For quantification, from both WT and OGT KO samples four 45 nm thin sections were imaged. For the eight composite virtual slides (one depicted in Figure 1 A) in total 7108 single 2k x 2k images were recorded containing 28.4 Gigapixel. Using a pixel size of 1.71 nm in total 8.3 10^4^ μm^2^ was imaged. Sections thickness was thinner that normal (90 - 100 nm) since these provided more structural details resulting in better models and predictions.

**Figure 1.**
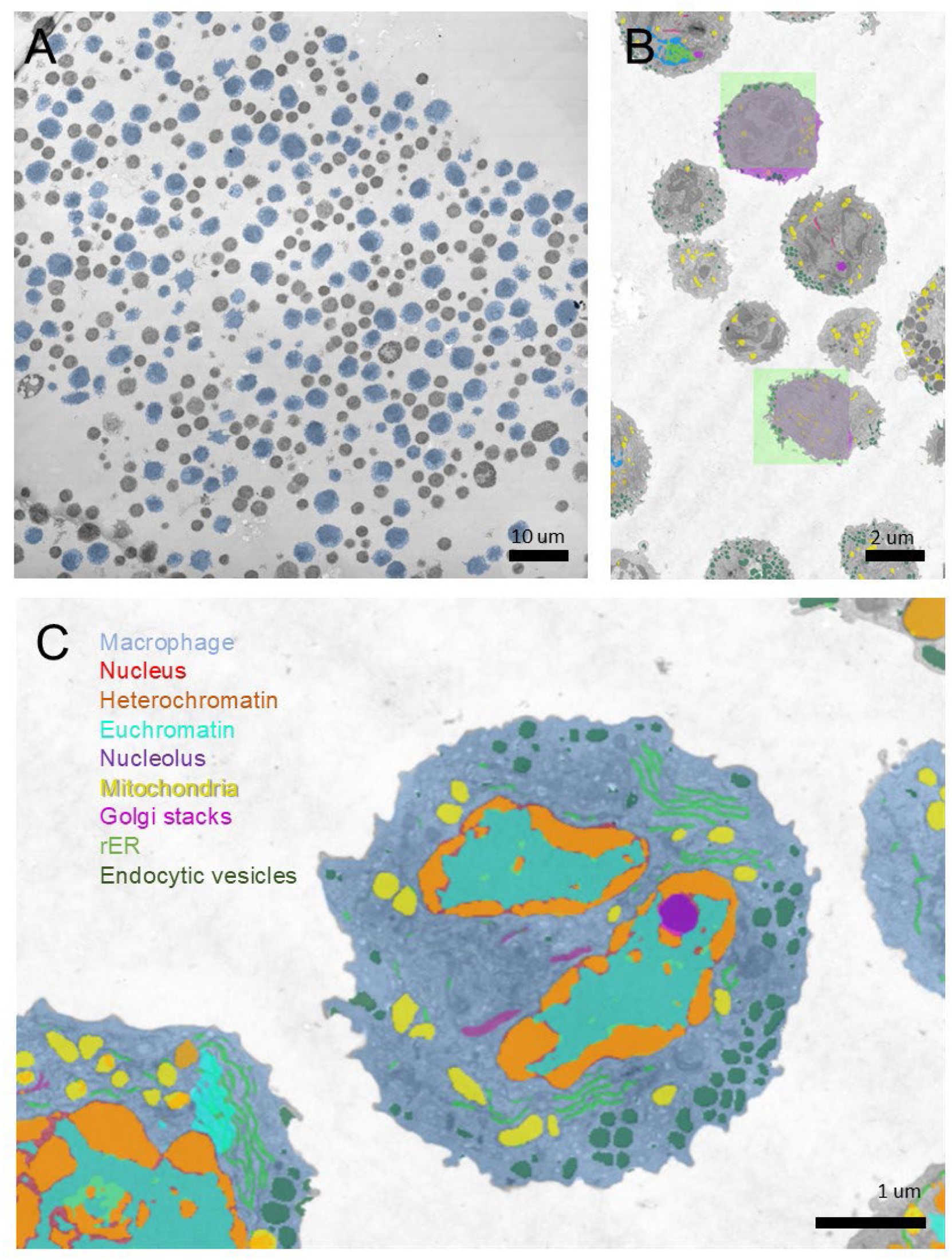
A) TEM overview image of section through cells harvested from the peritoneal murine cavity containing mainly macrophages (blue) and lymphocytes. B) Example image of manually annotated macrophages (purple) and boxes (green) used for CNN model generation and predicted, manually corrected macrophages (blue). C) Example of predicted subcellular structures inside a macrophage (blue), denoted are the nucleus (red), mostly overshadowed by heterochromatin (orange), euchromatin (cyan) and nucleolus (purple), mitochondria (yellow), rough ER (light green), endocytic vesicles (dark green) and Golgi stacks (dark pink).

Macrophages, nuclei, mitochondria, rough endoplasmic reticulum, Golgi stacks, endocytic vesicles, and within the nuclei heterochromatin, euchromatin and nucleoli were annotated by hand (and example of boxes and annotations are depicted in Figure 1B). Typically between 1 and 4% of the total image area was used for model building. For all structures CNN models were calculated and predictions were made, except Golgi stacks and nucleoli, which were completely segmented by hand. Predicted segmentations (see an example of one cell in Figure 1C) of macrophages and nuclei were corrected by hand, since it was important to have these correct since these formed the basis of the quantifications, while also for euchromatin minor corrections were performed. A total of 431 WT macrophages (from 2 mice) and 338 OGT KO macrophages (from 3 mice) were used for calculations.

### OGT KO macrophages are increased in size

From initial light microscopy observations it appeared that OGT KO mouse macrophages cells appeared enlarged compared to WT macrophages (Heieis et al., 2026; Yang et al., 2020). Since this was also apparent in the TEM stitches we first investigated differences in the macrophage size. Initially, the average macrophage size was calculated by dividing the total surface area of all macrophages by the number of cells, showing that OGT KO macrophages (64.8 ± 38.5 μm^2^ n=338) were ∼50% larger than WT macrophages (42.3 ± 19.3 μm^2^ n=431). We however suspected that these results did not correctly reflected the size difference. First, it appeared that not all macrophages from the conditional OGT KO mice displayed an equal loss of OGT activity, as evidenced by residual staining for the O-GlcNAc residue by flow cytometry (Heieis et al., 2026). Moreover, TEM sections through macrophages display a variable cell area size that is dependent on the position of the section through the cell, resulting in underestimation of the mean value of the sections compared to the actual size. To decrease the variance created by the sectioning bias and contamination of WT cells in the OGT KO sample, the sizes of the 88 biggest macrophages in both groups were determined (Figure 2A). The largest OGT KO macrophages have an average area of 117 ± 29.5 µm^2^ which is about twice as large as WT macrophages: 67.6 ± 8.8 µm^2^ (Figure 2B). From the images, the largest OGT KO cells furthermore appeared more oval compared to more circular WT macrophages (Figure 2A). Therefore, also the roundness of the 88 largest cells was determined. From violin plots (Figure 2C) macrophages appear to have a bimodal roundness distribution, suggesting two different states. OGT KO macrophages have a shifted distribution, especially of the more circular state, towards a more oval structure.

**Figure 2.**
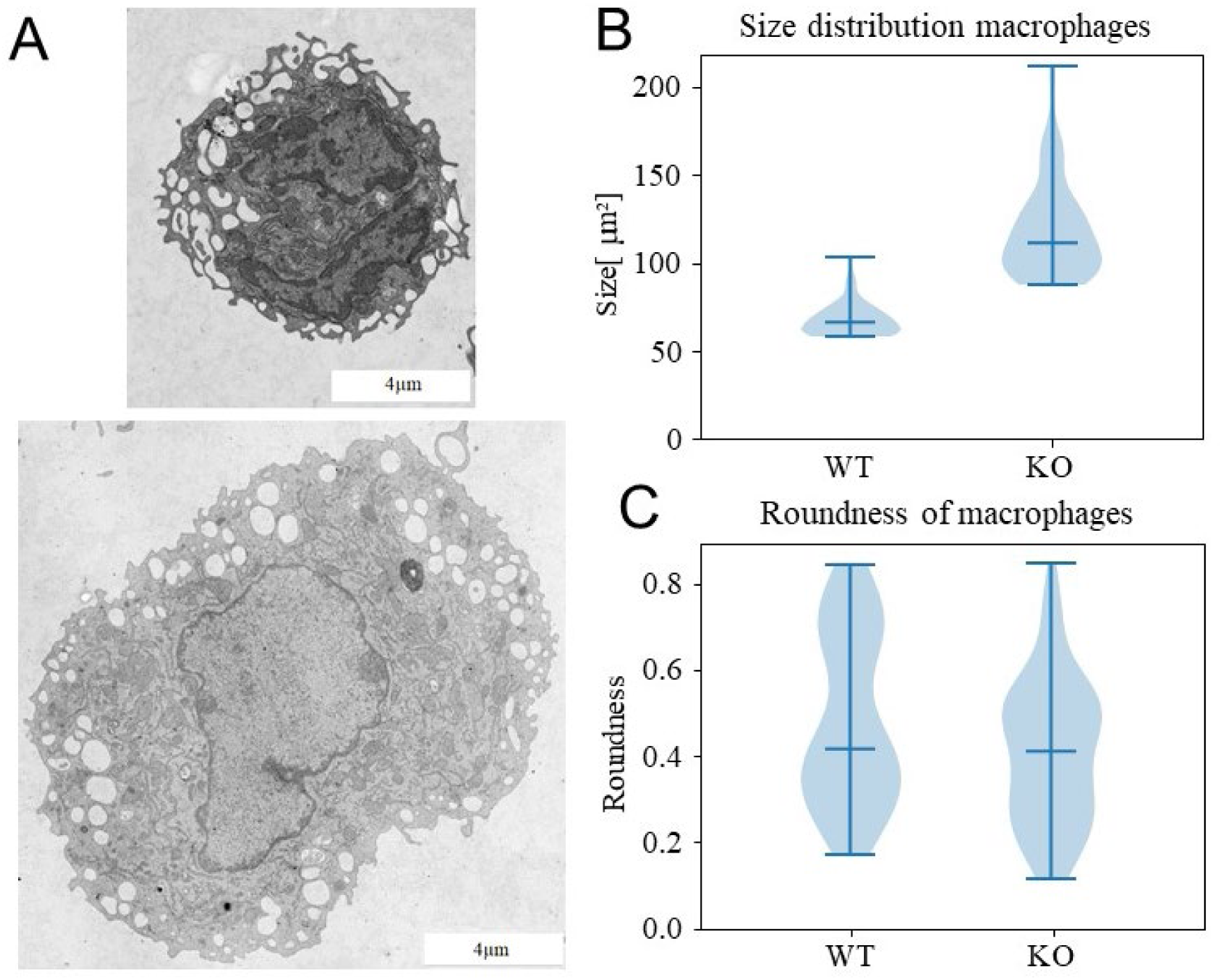
A) representative examples of WT (top) and OGT KO (bottom) macrophages with rough cytoplasmic membrane containing many cytoplasmic invaginations and vesicles. B) Cell size distribution of the largest 88 macrophages show that OGT KO cells are almost twice as large as WT cells. C) Roundness distribution measurements showing that OGT KO macrophages are more elongated than WT macrophages (n=88).

### Nuclear composition differs between WT and OGT KO macrophages

Given the increased size of the OGT KO macrophages and its potential causes, we analysed the effects of OGT deficiency on the nuclei and its in EM sections structurally discernible components, heterochromatin, euchromatin and the nucleolus.

Analysis of the segmented nuclei showed that the absolute size of nuclei from OGT KO macrophages (12.0 ± 8.6 µm^2^ n=290) was roughly 50% larger than nuclei from WT macrophages (8.5 ± 6.2 µm^2^ n=339) (Figure 3A) and that the nuclei from OGT KO macrophages have a more bimodal distribution that the nuclei from WT macrophages. Since OGT KO macrophages themselves are also larger, also the relative size differences of the nuclei compared to the macrophages were evaluated. The relative nucleus size in OGT KO macrophages did not significantly differ from WT macrophages (Figure 3B), meaning that the nucleus size increases at the same rate as the cell size, but does not account for the full increased cell size.

**Figure 3.**
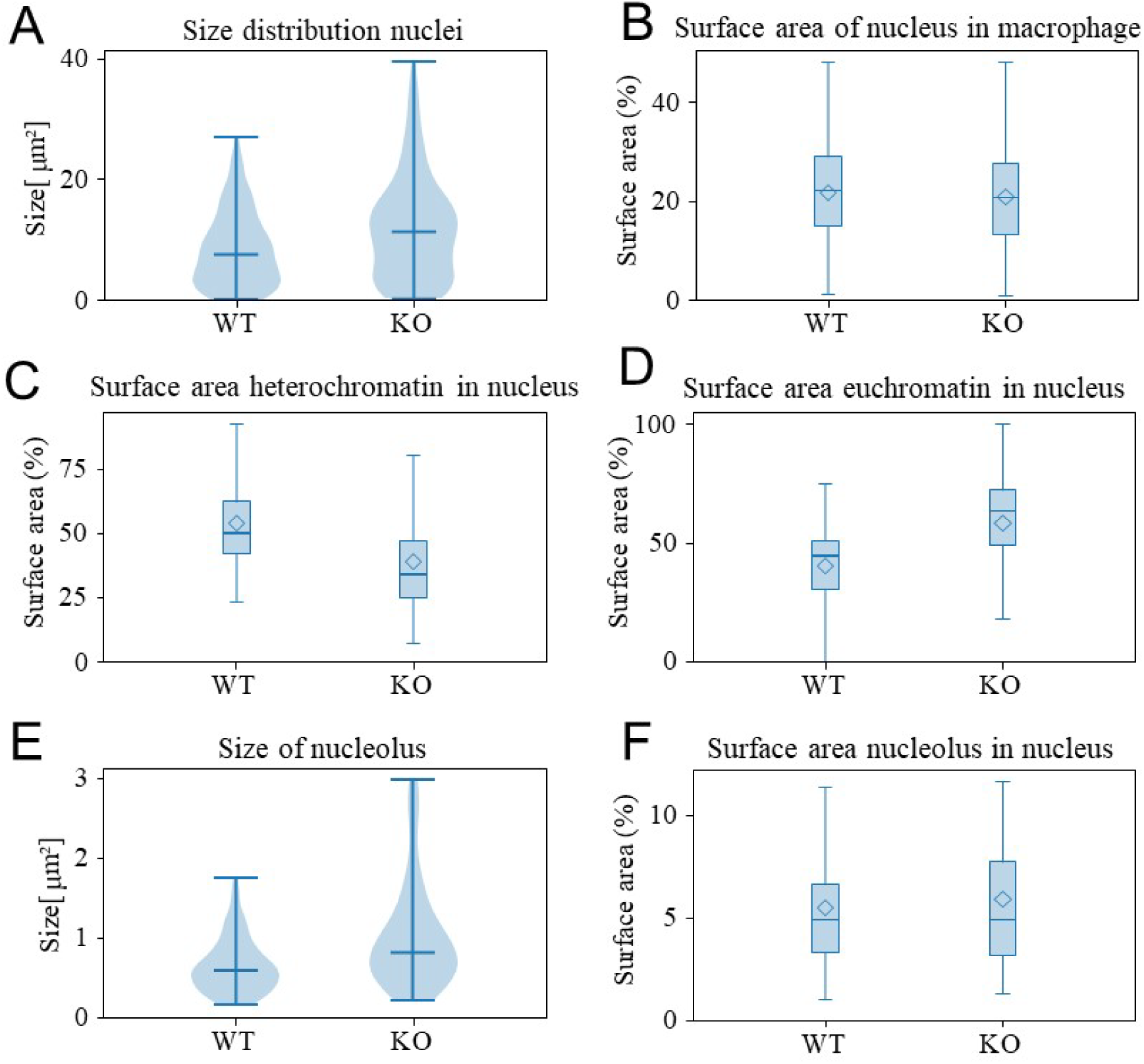
A) Size distribution of nuclei in WT (n=300) and OGT KO macrophages (n=220). Surface area percentage of WT and OGT KO macrophages B) nuclei in the whole cell, and C) heterochromatin and D) euchromatin in nucleus. E) Size distribution of nucleoli in WT (n=126) and OGT KO (n=111) nuclei. F) The surface area percentage of the nucleolus in the nucleus. Data are median (center line) ± SEM and mean (diamond).

Next, from heterochromatin, euchromatin, and the nucleolus the relative the surface areas to the nucleus were compared. From visual observations there appeared to be more euchromatin (lighter areas) in OGT KO macrophages compared to WT macrophages (Figure 2A), which was confirmed by quantification, which showed that the surface area euchromatin in nuclei in OGT KO macrophages is roughly 50% (58.6 ± 20.0 %, n=220) than in nuclei of WT macrophages (40.5 ± 15.9 %, n=300) (Figure 3C, D). Similarly, the nucleoli in OGT KO macrophages (1.0 ± 0.66 µm^2^, n=111) had a slight increase in absolute size compared to nuclei from WT macrophages (0.64 ± 0.34 µm^2^, n=126) (Figure 3E), while their relative size to the nucleus was similar (5.5 ± 3.5 % in WT vs. 5.9 ± 3.9 % in KO) (Figure 3F).

### rER and stacked Golgi amount have increased in OGT KO macrophages

The observed increased amount of euchromatin, decreased amount of heterochromatin, and a larger nucleolus, are consistent with an increase of DNA transcription in OGT KO macrophages. Therefore, organelles involved in the downstream protein synthesis might also experience changes in number or size, due to upregulated protein synthesis and general cell activity. So next, we analysed rER and Golgi apparatus, which regulate protein synthesis, folding, transport and post-translational modifications. Our results indeed reflected an increase in surface area of these organelles. The relative surface area of rER in OGT KO macrophages (3.5 ± 1.8%, n=24321) was higher than in WT macrophages (2.6 ± 1.4%, n=16982) (Figure 4A). For the Golgi apparatus stacks, a slight increase in surface area of the Golgi stacks was observed in OGT KO macrophages (0.8 ± 0.6%, n=319) compared to WT macrophages (0.6 ± 0.4%, n=379) (Figure 4B). Of note, the actual size of the Golgi apparatus includes small vesicles and is larger than just the Golgi stacks that were manually annotated and quantified.

**Figure 4.**
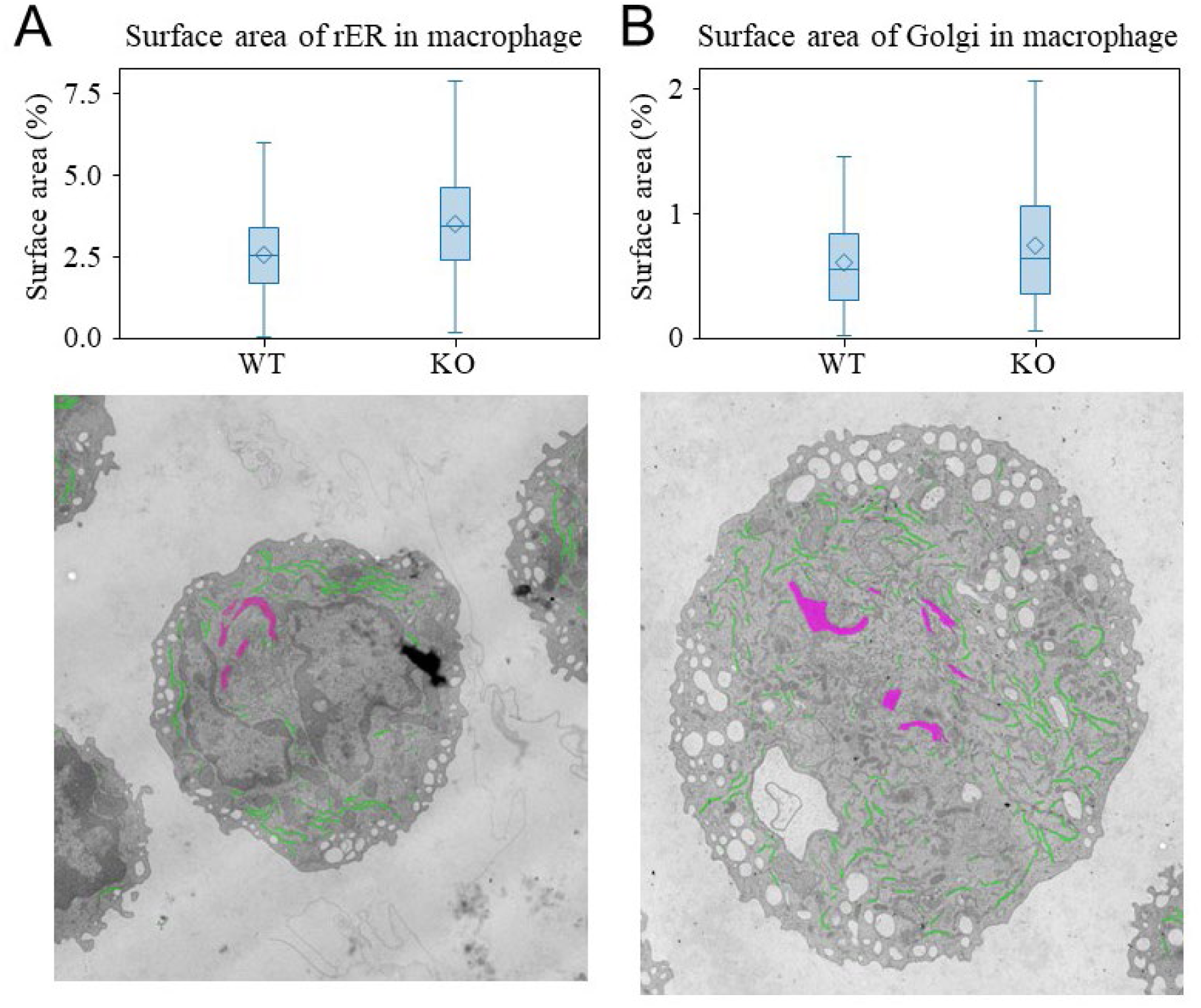
Surface area percentage distribution of A) rER and B) Golgi apparatus in the cell. Boxplots show median (line) ± SEM and mean (diamond). Example images of C) WT and D) OGT KO macrophages.

### Mitochondrial shape and size are not equal in WT and OGT KO macrophages

Increased presence of euchromatin and, rER and Golgi might suggest increased cellular activity requiring increased ATP consumption and changes in mitochondrial activity. Therefore, we analysed mitochondrial occurrence, size, relative presence, and shape. In total 5284 mitochondria from 431 WT macrophages and 6951 mitochondria from 328 OGT KO macrophages were analysed. The absolute number of mitochondria per macrophage differed between the groups. In OGT KO macrophages 50% more mitochondria (21 ± 15) were observed in the cellular sections per macrophage than in WT cells (14 ± 9) (Figure 5A). Also, the mitochondrial surface area within OGT KO macrophages (3.9 ±2.3%; n=431) was higher compared to WT macrophages (2.9 ± 1.9%; n=328) (Figure 5B).

**Figure 5.**
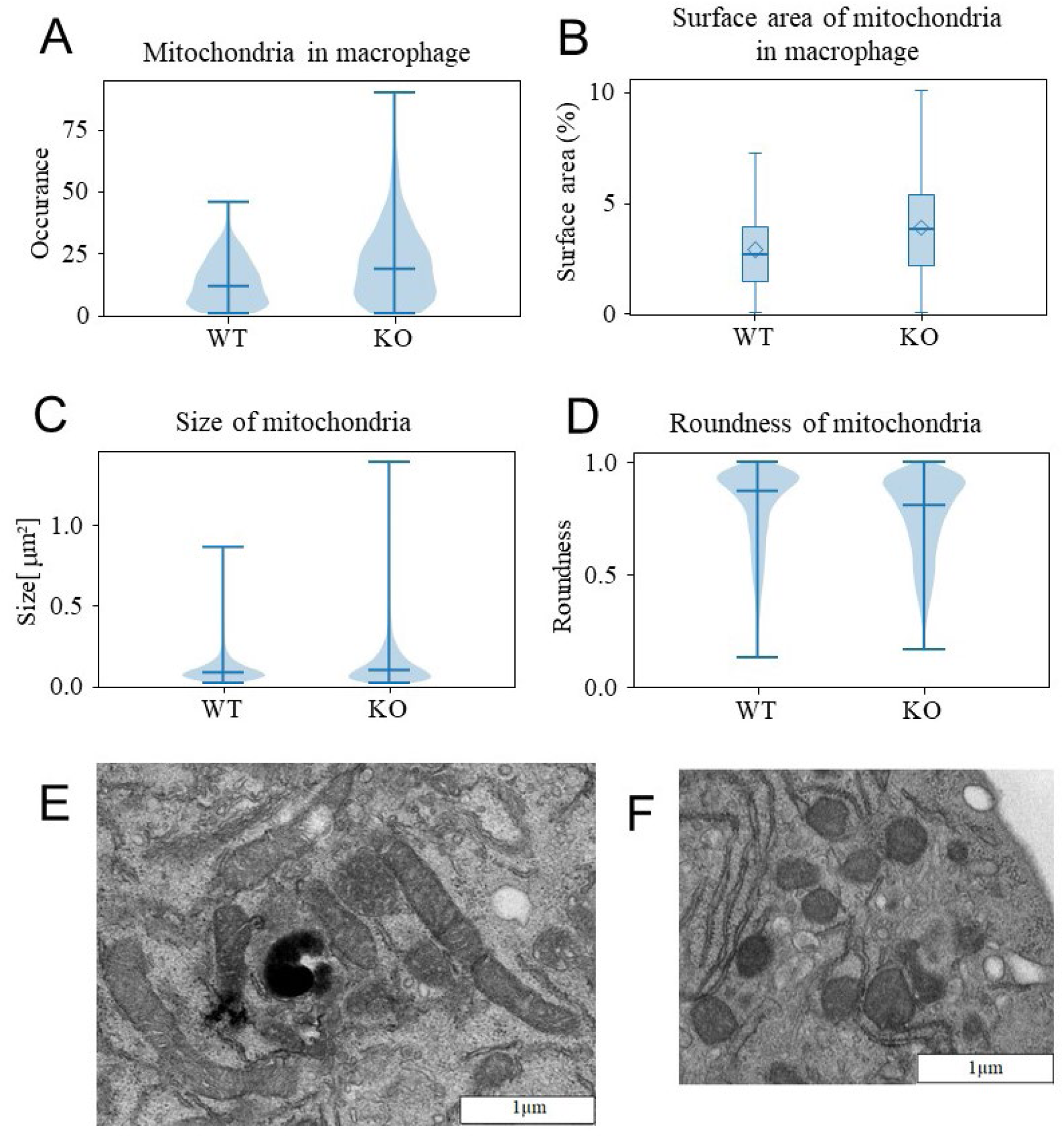
Mitochondria are more abundant and larger in OGT KO macrophages A) Violin plot showing the occurrence distribution of mitochondria in WT (n=431) and OGT KO (n=338) macrophages. B) The surface area percentage of the mitochondria in macrophages. C) Violin plot of mitochondria size distribution in WT (n=5284) and OGT KO (n=6951) macrophages. D) Violin plot showing the roundness distribution of mitochondria in WT and OGT KO macrophages. E) TEM image of OGT KO macrophages shows that mitochondria are elongated and can form networks, while F) mitochondria in WT macrophages appear rounder and smaller. Violin plots show median (center line) ± SEM and mean (diamond).

The mitochondria in OGT KO cells were also slightly larger (0.12 µm^2^ in OGT KO vs. 0.10 µm^2^ in WT) (Figure 5C) and longer (roundness: 0.76 ± 0.18 in OGT KO vs. 0.81 ± 0.17 in WT) (Figure 5D), indicating that mitochondria in OGT KO macrophages tend to form more elongated tubular structures, which could also be observed in the TEM images (Figure 5E, F). Furthermore, close observation of the TEM images also showed that the outer membrane of the mitochondria had a more wrinkled appearance, while WT mitochondrial surfaces were smoother.

### Endocytic vesicle quantity is equal in both cell types

To conclude, we analysed the occurrence, size, relative presence, and shape of the endocytic vesicles. A major function of macrophages is to clean up the remains of dead cells and foreign particles, by engulfing them with the cell membrane into inclusion or endocytic vesicles, where these objects are digested. Because OGT KO macrophages are increased in size, they have a larger membranous area, and it could be argued that this affects the number of endocytic vesicles. Our findings suggest that this is not the case. On average, the OGT KO macrophages possessed similar numbers of endocytic vesicles per cell section as WT macrophages (46 ± 39 in KO vs. 48 ± 27 in WT) (Figure 6A), which, on average were slightly larger (0.13 ± 0.11 µm^2^ in KO vs. 0.10 ± 0.10 µm^2^ in WT) (Figure 6B) and similar in shape (roundness: 0.84 ± 0.16 in KO vs. 0.85 ± 0.16 for WT) (Figure 6C). So the size, occurrence and shape of the endocytic vesicles did not significantly differ between OGT KO and WT macrophages.

**Figure 6.**
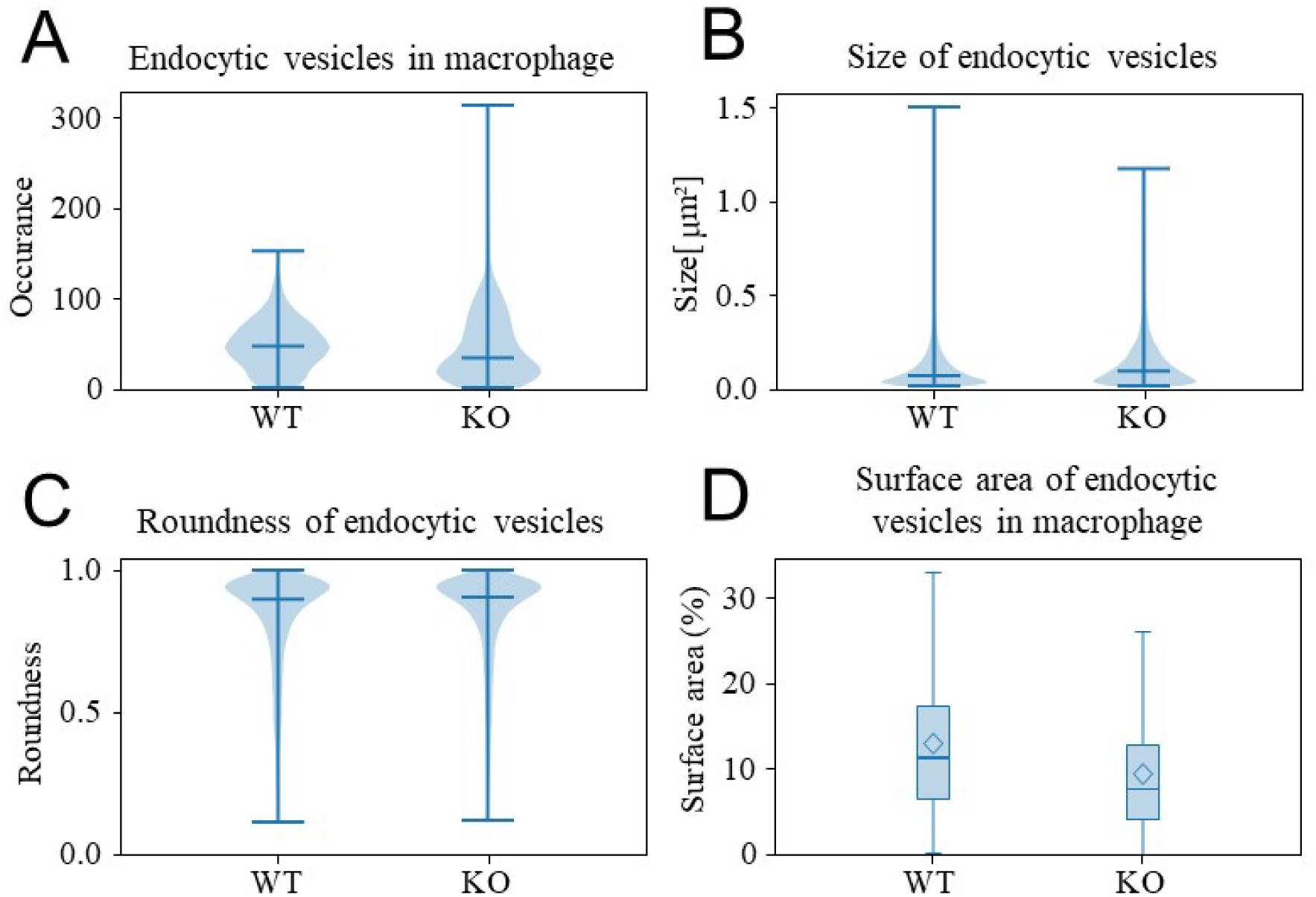
Endocytic vesicles are less abundant in OGT KO macrophages A) The number of endocytic vesicles per cell has decreased in OGT-deficient macrophages. B) The size of the endocytic vesicles increases in OGT KO macrophages. (C) Violin plots of the roundness of endocytic vesicles shows that their shape is similar in both WT and OGT KO macrophages. D) Overall, the endocytic vesicles (n=20827) in WT cells have a larger surface area percentage than endocytic vesicles (n=15575).

### TEM expert image assessment

Two important remaining questions following machine learning segmentation and quantification of TEM slides of plastic embedded and stained macrophage sections are (i) whether the results fit biological functional changes of the OGT KO macrophages, and (ii) whether the results agree with more conventional way of analysis of the TEM images by human experts. Comparison of the functional implication of the TEM analysis are analysed in-depth and described in a separate manuscript (Heieis et al., 2026). Here, we focus on the analysis by six experts and the comparison with the sML quantification (Table 1).

**Table 1.**
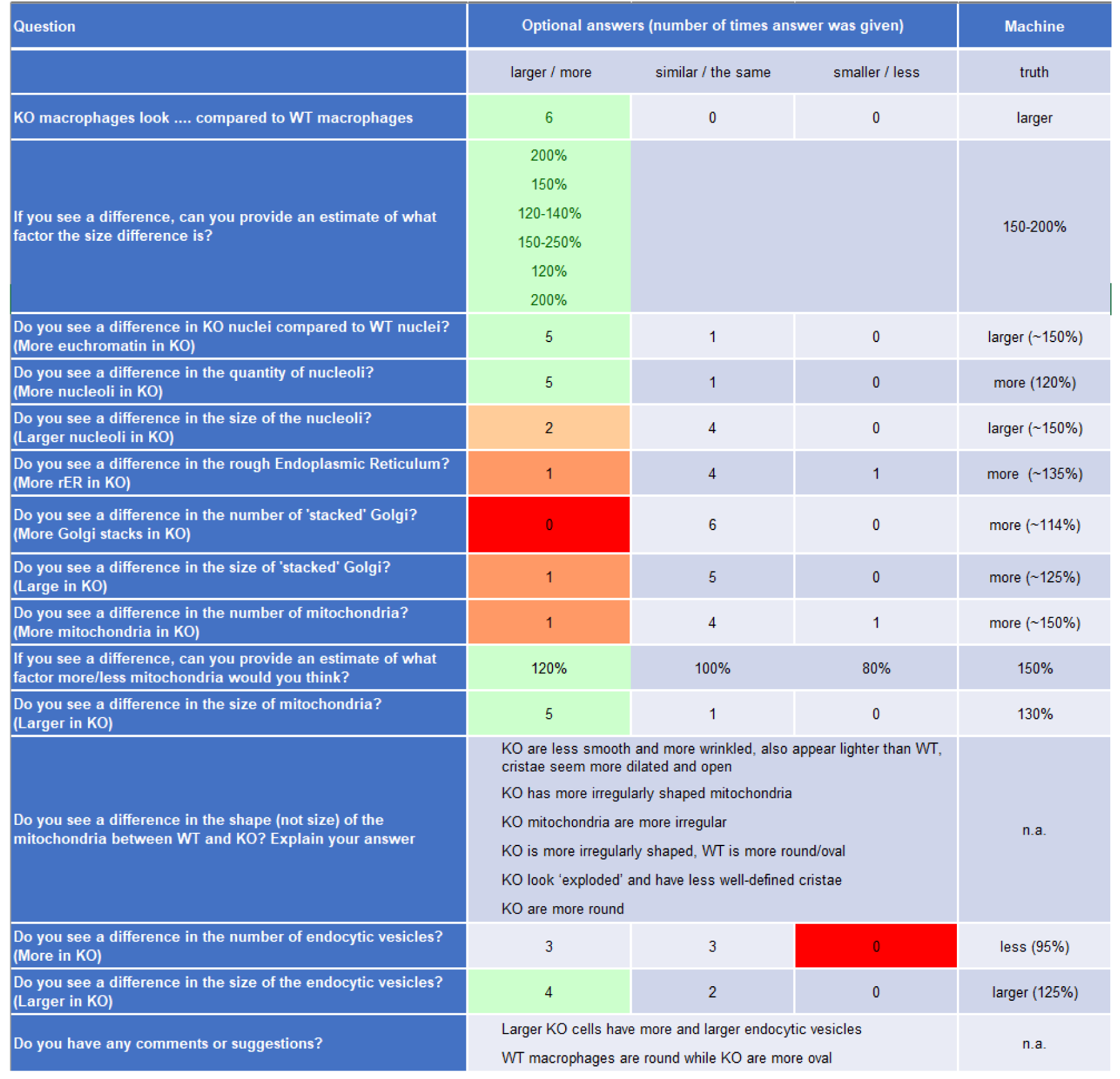
a Overview of questionnaire and answers from six TEM experts compared to machine earning results. Agreement of TEM expert and machine learning quantification results are colour coded from green (good match) through orange (reasonable match) to red (bad match).

All experts spotted the size difference of WT and OGT KO macrophages. The size differences were estimated quite well, varying from 120% to 250%, which was in accordance with the calculated size increase of all cells (∼150%) and the largest cells only (∼200%) using sML. Without specifically asking for it, two out of six experts also noted that OGT KO macrophages appeared more oval. Increased size of the nucleus and presence of euchromatin was spotted by 5 out of 6 experts. Also, morphological differences of the mitochondria, including size differences, were well detected (5/6). Interestingly, differences in occurrence and size of both sparsely present structures, such as the nucleoli, and the stacked Golgi, as well as extensively present and widely distributed structures, such as the rER and mitochondria, were missed by the experts in 80% of the cases. Very small changes, 14% change in Golgi stack size and 5% change in number of endocytic vesicles, were completely missed by all experts.

## Discussion

The goals of this study were to evaluate the potential of supervised machine learning as a tool for segmentation, quantitation, and interpretation of virtual TEM slides to obtain a detailed quantitative understanding of ultrastructural changes in macrophages following OGT KO, in order to. In particular, we addressed the question of whether AI-based quantitative analysis can complement or even replace conventional qualitative interpretation by experienced TEM experts.

Initial biochemical characterization of OGT-KO macrophages indicated substantial changes in their cell size, mitochondrial polarization, transcriptional activity and protein translation. To link these biochemical changes to morphological variations we performed systematic segmention and quantification TEM stitches, assessing changes in occurrence, relative surface area, size and shape of cellular ultrastructure. The biological implications of these morphological parameters in relation to macrophage cellular processes and function are discussed in detail elsewhere (Heieis et al., 2026). Here we primarily focused on the methodological aspects of machine learning assisted quantitative image analysis and compared its performance to the current benchmark (i.e. conventional and qualitative human TEM expert interpretation).

### Supervised machine learning for ultrastructural segmentations

To achieve large-scale quantitative analysis, we used supervised machine learning to segment a wide range of different organelles an structures in composite 2D TEM images. While sML has previously been shown to perform well in 3D scanning electron microscopy datasets (Xiao et al., 2018), its application to large stitched TEM images presents distinct challenges due to image size, structural complexity, and variability in contrast. Using a custom-built web interface incorporating a TensorFlow/Keras-based convolutional neural network (Dzyubachyk et al., 2021) adapted for large 2D TEM composite images (Faas et al., 2012), we generated predictive models and predictions for multiple ultrastructural components. Our results demonstrate that sML segmentation is feasible and reliable for subcellular structures with distinctive morphology and sufficient representation in the training data, such as nuclei, heterochromatin, euchromatin, mitochondria, and endocytic vesicles. These models enabled the analysis of a statistically substantial number of cells at high spatial resolution, which would be impractical using full manual annotation alone.

### Methodological limitations of 2D TEM-based quantification

Interpretation of quantitative TEM data must account for the inherent limitations of two-dimensional sections through three-dimensional cells. To obtain representative measurements, both equal probability of sectioning and unbiased probing are essential (Boyce, Dorph-Petersen, Lyck, & Gundersen, 2010). In our macrophage suspension, cells were free-floating prior to embedding, ensuring random orientation and equal likelihood of sectioning through any cellular region. However, unbiased probing remains a constraint due to section thickness. With cells approximately 10 µm in diameter and section thicknesses of 45–90 nm, even relatively large datasets correspond to only a small fraction of whole-cell volume. When all sections are combined, the total analyzed area effectively represents approximately one whole cell per condition. This highlights the necessity of large datasets and systematic analysis to reduce sampling bias—an area where automated segmentation provides a clear advantage over manual approaches.

### Ultrastructural consequences of OGT knockout in macrophages

Quantitative analysis revealed a clear increase in macrophage size following loss of OGT, with average diameters increasing from 9.2 µm in WT to 12 µm in OGT KO cells. To obtain reliable size estimates, only the largest 88 cells from each group were analysed. This approach corrects for two confounding factors: incomplete knockout efficiency (with an estimated 10–20% residual WT macrophages) and the geometric bias inherent to random thin-section sampling of spherical cells, where smaller profiles often represent off-centre sections.

The measured WT macrophage size was smaller than values previously reported for rat macrophages (13.1 µm) (Krombach et al., 1997), likely reflecting species differences and methodological variation (TEM vs. flow cytometry). The observed increase in cell size and altered morphology are consistent with changes in macrophage activation state (McWhorter, Wang, Nguyen, Chung, & Liu, 2013).

Although nuclear size was higher in absolute terms in OGT KO macrophages, the nuclear-to-cell surface ratio remained constant, indicating proportional growth of the nucleus with the cell. The nuclear enlargement was driven primarily by increased euchromatin content and enlarged nucleoli, suggesting elevated transcriptional and translational activity. These findings align with a more metabolically active macrophage phenotype (McWhorter et al., 2013).

Consistent with this interpretation, we observed modest increases in rough endoplasmic reticulum (rER) and Golgi stack surface area in OGT KO macrophages. It should be noted that only Golgi membrane stacks were segmented, excluding surrounding vesicles, and that rER segmentation was technically challenging due to morphological variability and overlap with other structures. Nonetheless, analyses using multiple rER models of varying quality yielded consistent trends, supporting the robustness of the observed increase.

Mitochondrial morphology was also significantly altered. Previous work demonstrated that O-GlcNAcase (OGA)-deficient fibroblasts exhibit smaller, more fragmented mitochondria (Akinbiyi et al., 2021). This was associated with increased O-GlcNAcylation of mitochondrial fission promoting factor Drp1. Given the antagonistic relationship between OGA and OGT, the opposite phenotype was anticipated in OGT KO cells. Indeed, mitochondria in OGT KO macrophages were larger (0.12 µm² vs. 0.10 µm² in WT) and more elongated, which are commonly linked to increased ATP synthesis. These changes likely reflect increased energetic demand in a more active cellular state. Additionally, the wrinkled appearance of the outer mitochondrial membrane suggests altered mitochondrial dynamics or function.

### sML vs. TEM expert interpretation

A central aim of this study was to compare objective, sML-derived quantifications with subjective assessments by six experienced TEM experts. Expert observations showed limited consistency: apart from the increase in cell size, no single ultrastructural change was unanimously identified. Human estimated size increases (120-250%) varied but were broadly consistent with quantitative measurements (150-200% for the largest cells), particularly given natural biological variability and sectioning effects.

Overall, 55% of the TEM expert observations did not align with the quantitative results, highlighting the difficulty of reliably estimating numerical differences in complex ultrastructural datasets by visual inspection alone. Notably, the magnitude of a quantitative differences of structures as predicted by sML, were not reflected in the positive detection by human experts. While a ∼50% increase in euchromatin area was recognized by most experts (5/6), similar increases in nucleolar size and mitochondrial number were rarely noted. Conversely, larger changes—such as the ∼125% increase in endocytic vesicle size and ∼130% increase in mitochondrial size—were more consistently identified.

In contrast, experts readily recognized subtle qualitative structural differences, including mitochondrial elongation, and irregularities of the outer mitochondrial membrane. These subtle shape and texture features are difficult to capture in current quantitative segmentation frameworks but are intuitively recognized by trained observers.

### Can AI replace human TEM expertise?

Our results show that sML outperforms human observers in the objective quantification of structural numbers, sizes, and surface areas across large datasets. However, AI cannot yet replace human expertise in ultrastructural TEM analysis. Experts remain essential for generating training annotations, manually segmenting rare or ambiguous structures, correcting prediction errors, and interpreting subtle morphological features.

## Conclusion

Large-scale imaging combined with machine-learning-assisted segmentation enables comprehensive, statistically analysis that is unattainable through manual inspection alone. Given the inherent biological variability of ultrastructure, the limitations of 2D sampling of 3D cells, and the subjective nature of human annotation, reliance on qualitative assessment alone carries a substantial risk of mis-or over-interpretation and should always be used in conjunction with orthogonal data.

We conclude that supervised machine learning is a valuable tool for quantitative analysis of large-scale TEM images, which is best used in close synergy with expert interpretation. Together, they enable a more objective, reproducible, and biologically meaningful understanding of ultrastructural changes.

## Acknowledgements

We would like to thank Dr. Carolina Jost, Dr. Mieke Mommaas, Cristina Avramut, Ronald W.A.L. Limpens, Aat A. Mulder for taking part in the TEM expert survey.

